# High-Activity Enhancer Generation based on Feedback GAN with Domain Constraint and Curriculum Learning

**DOI:** 10.1101/2023.12.16.570150

**Authors:** Jiahao Li, Liwei Xiao, Jiawei Luo, Xianliang Liu, Junjie Chen

## Abstract

Enhancers are important cis-regulatory elements, enhancing the transcription of target genes. De novo design of high-activity enhancers is one of long-standing goals in generated biology for both clinical purpose and artificial life, because of their vital roles on regulation of cell development, differentiation, and apoptosis. But designing the enhancers with specific properties remains challenging, primarily due to the unclear understanding of enhancer regulatory codes. Here, we propose an AI-driven enhancer design method, named Enhancer-GAN, to generate high-activity enhancer sequences. Enhancer-GAN is firstly pre-trained on a large enhancer dataset that contains both low-activity and high-activity enhancers, and then is optimized to generate high-activity enhancers with feedback-loop mechanism. Domain constraint and curriculum learning were introduced into Enhancer-GAN to alleviate the noise from feedback loop and accelerate the training convergence. Experimental results on benchmark datasets demonstrate that the activity of generated enhancers is significantly higher than ones in benchmark dataset. Besides, we find 10 new motifs from generated high-activity enhancers. These results demonstrate Enhancer-GAN is promising to generate and optimize bio-sequences with desired properties.

## I. Introduction

Enhancers exert cis-regulation over target gene expression, thereby influencing biological development and physiology [1–3]. Enhancer generation is an important but challenging task in generated biology. Nonetheless, the challenge of designing and optimizing novel enhancers with desired properties persists mainly due to the vague understanding of enhancer regulatory code and their high dimensionality [4]. Current technologies for designing enhancers with desired properties are mostly manual and require significant expert experience, such as mutagenesis and recombination of motifs [5], but these approaches were confined to a relatively narrow sequence space, preventing the exploration of improved sequences in a broader sequence space, despite the latter option imposing a significant burden on biological experiments.

AI-based generative models, such as generative adversarial networks (GANs), can learn the biological motifs from extensive enhancer sequences and generate novel enhancers meeting the required properties. Approaches of using GAN for generated biology undergo unsupervised training to approximate the distribution of real data and then generate the novel sequences. However, GANs often encounter issues like mode collapse and struggle with generating heavy tail distributions, especially in specific biology applications with spare data, which amplifies the challenge producing the meaningful samples. Feedback GAN [6] is promising to address these issues by incorporating a feedback-loop mechanism, but it suffers from the noise of the external analyzer and slow speed in training. Because the generalization of external analyzer usually lacks comprehensive evaluation on random sequences and unknown sequences. Thus, the feedback-loop mechanism can’t provide promising feedback when there are no constraints on generated sequences, which could lead to the generative models failing to produce high-quality sequences.

In this paper, we present a novel feedback GAN with domain constraint and curriculum learning, called Enhancer-GAN, for high-activity enhancer generation. Enhancer-GAN is firstly pre-trained on the a large enhancer dataset to learn the general grammar of enhancer sequences, and then is optimized with a feedback-loop for high-activity enhancers (**Fig. 1**). During feedback-loop training, low-activity enhancers in training dataset will be replaced with high-activity generated enhancers when their sequence features satisfying the domain constraint and their activity above threshold in curriculum learning. Our main contributions are summarized as follows:

1. We introduce the domain constraint into the feedback mechanism to alleviate the noise from the external analyzer when optimizing the generated sequences for high activity.
2. We adjust the activity threshold used in feedback-loop by the curriculum learning according to the current generated sequences, which will speed up the model convergence. 3) Enhancer-GAN enable us to extract vital motifs from high-activity enhancers, which is helpful to understand the biological mechanism of enhancers.

**Fig. 1:**
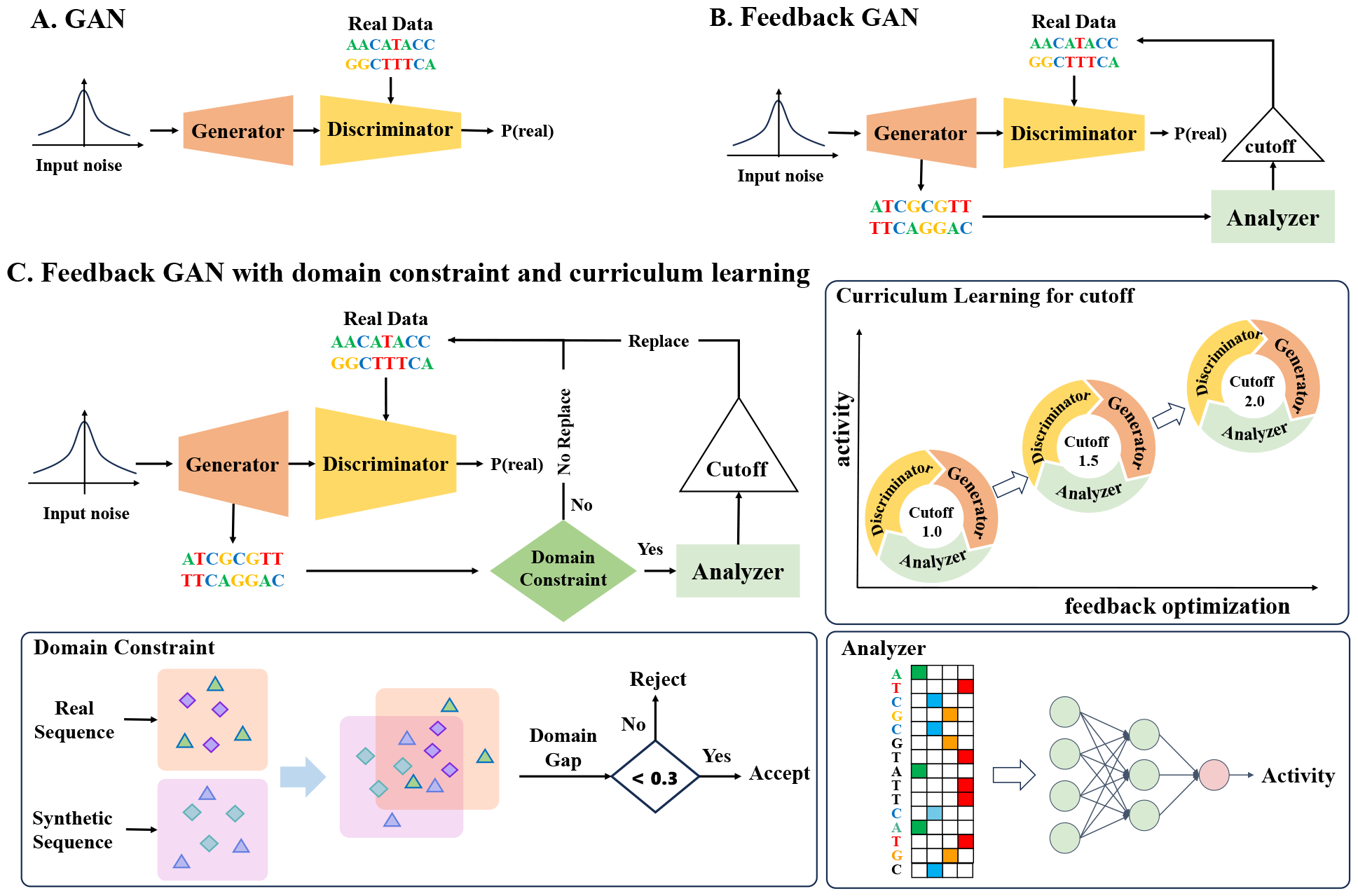
The framework of Enhancer-GAN with domain constraint and curriculum learning for high-activity enhancer generation.

## II. Related Work

Recent advances in deep learning methods have supported human creativity for biology sequences, including molecular design [7, 8], protein generation [6, 9] and regulatory DNA [10], often with desired properties.

These methods can be grouped into three categories: conditional approaches with label information, post-screening on generated sequences and the optimization approaches with external predictors. Conditional approaches integrate label information into the generation model to produce sequences with desired properties [11], but it is limited by the available data between sequences and conditional information. Post-screening based methods first generate a large number of candidate sequences and then screen sequences that meet the required properties by a filter [12]. Optimization approaches use external predictors to guide the design to target properties through the iterative training-generation-prediction steps, without any consideration of gradients from the external predictors. For example, Hoffman et al.[13] optimized the decoder for desired properties based on zeroth-order optimization. Gupta et al.[6] incorporated the feedback-loop mechanism for the optimization. Mokaya et al.[14] optimized generated molecules for desired profiles based on the curriculum learning.

These existing methods rely on an external analyzer for property evaluation in their optimization, while ignoring its generalization due to the lack of comprehensive evaluation on random sequences and unknown sequences. Thus, they suffer from the noise from the external analyzer when there are no constraints during optimization, thereby misleading the generator to produce low-quality sequences. So we introduce the domain constraint and curriculum learning in feedback GAN to optimize generated enhancers with high activity.

## III. Method

### A. Dataset and metric

To evaluate the proposed method, we selected enhancers from [3] as the benchmark dataset, containing 242,014 enhancer sequences extracted from the Drosophila S2 genome by UMI-STARR-seq. All enhancer sequences are in the same length of 249bp, with the average activity of 0.37, where 48% of sequences have activity less than 0.0, just 2% higher than 3.0. The dataset comprises three sub-datasets, containing 201,138, 20,592, 20,284 enhancer sequences, respectively. These subsets are employed in the stages of pre-training, feedback optimization and validation, respectively.

We employ Maximum Mean Discrepancy (MMD) [11] as the metric to evaluate the quality of generated enhancers. The formula for MMD is shown as Eq. (1).

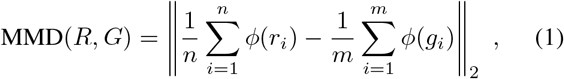

where 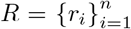 and 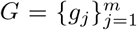 are sampled from the distributions of real and generated enhancers, respectively.

To calculate the MMD of generated and real enhancers, we need to first project all enhancers into a high-dimension representation space. In this paper, we adopt two embedding methods: the normalized Spectrum kernel[15] and DNABert [16]. And finally, we chose *k*mer with 3,4,5,6 for both methods considering the trade-off between vector dimension and information of each sequence.

### B. Generative adversarial networks

In our work, we employed WGAN as our framework (**Fig. 1A**), which employs the wasserstein distance to mitigate model collapse in vanilla ones [17, 18]. WGAN was pre-trained to capture the general grammars of enhancers. We introduced a feedback-loop mechanism [6] to optimize the WGAN model for generating high-activity enhancers. The generator plays the same game with the discriminator as usual, but additionally generated samples will be sent to an external analyzer to estimate their biological activity (**Fig. 1B**). These generated enhancers with activity scores above the cutoff are selected to replace the original enhancers with low activity in the discriminator’s training dataset. In the feedback loop, the training dataset of the discriminator is gradually replaced by high-activity generated sequences. Therefore, the generator can focus more on the high-activity sequences in the mini-max game with the discriminator.

The analyzer we used in feedback-loop mechanism is a well-trained model, DeepSTARR[3], with a outstanding performance of PCC=0.68 (**P**earson **C**orrelation **C**oefficient). It consists of four one-dimensional convolution layers and two fully connected layers, which directly takes DNA sequences as input and then quantitatively predicts their activities.

### C. Feedback mechanism with domain constraint and curriculum learning

The analyzer may not deliver robustness feedback when encountering generated samples beyond its generalization capacity, potentially introducing false positive samples into the optimization and leading to mode collapse. Besides, the preset activity threshold, *cutoff*, is a vital parameter, which determines the quality of feedback from the analyzer. A high value will increase the challenge of model optimization, while a low value limits the model optimization to learn the distribution of high-activity enhancers. To end these issues, we introduced the domain constraint [19] and curriculum learning [20] in the feedback loop (**Fig. 1C**).

#### Domain Constraint

To alleviate noise from the analyzer, we added a domain constraint into the feedback loop to reduce the false positive sequences. The idea of domain constraint is intuitive. Since the analyzer is trained on the finite number of enhancers, its generalization bound is not evaluated on unseen sequences. To make analyzer provide promising feedback, its inputs are constrained in a similar domain with the training dataset of analyzer. In this paper, we utilized MMD to measure the domain gap between two distributions. The replacement in feedback loop will be skipped if the *gap* between generated sequences and validation sequences is above a threshold *T*_*gap*_. Regarding the trade-off between quality and high-activity of generated enhancer, we set *T*_*gap*_ as 0.3 in our experiments.

#### Curriculum Learning

Curriculum learning is to start with similar and easier sub-tasks, and then gradually increase the difficulty level until finishing the whole task. The goal of pretrained GAN is to learn general grammar of enhancers that are usually low activity, while the goal of feedback mechanism is to optimize the GAN model for high-activity enhancers.

We used curriculum learning to guide this transformation, gradually improving the goal of optimization from an easy task of generating low-activity enhancers to a complex task of generating high-activity enhancers. Specifically, we gradually raised the cutoff in the feedback mechanism to control the difficulty of transformation. It was set relatively low at the beginning, and raised gradually when the model can generate valid enhancers and their average activity exceeds cutoff.

### D. Motifs extraction

Enhancers consist of binding sites (motifs) that control gene expression [21]. Here, we extracted motifs from the generated enhancers during the feedback optimization process to explore the relationship between motifs and regulatory activity.

We saved 500 intermediate models during the optimization of pre-trained model in the feedback with curriculum learning. We fed the same noise into the 500 saved models and then collected the corresponding 500 generated enhancers, named as **evolution set**, because such the process is analogous to the evolution of a raw enhancer from low-activity to high-activity. To find motifs enriched in the high-activity enhancers, all generated enhancers are divided into three groups according to their activity, high-activity group with activity larger than 3.0, low-activity group with activity less than 0.0 and mid-activity group with activity between them. STREME [22, 23] was employed to find relatively enriched motifs in each group.

## IV. Experiments & Results

### A. Experimental setup

Both the generator and the discriminator in WGAN were built with five residual blocks, and each residual block consists of two one-dimensional convolutional networks and activation functions of ReLU, without any use of normalization. In addition, multi-layer perceptron (MLP) following residual blocks was to translate the output of the last residual block into the target results, where the MLP in the generator outputs the generated enhancers and that of the discriminator discerns the source of input samples. All layers were initialized by the algorithm of Xavier-Uniform. The ratio of G and D training steps was set as 1:5. And the Adam algorithm (*β*_1_ = 0.5 and *β*_2_ = 0.9) was used. All models were trained with Nvidia GEFORCE RTX 4090 and Intel Xeon Platinum 8358P.

The training process of Enhancer-GAN contains two phases: pre-training and optimization. The pre-training phase was trained on the large pre-trained sub-dataset (201,138 enhancers with the average activity of 0.37) to learn the general grammar of enhancers (**Fig. 1A**). The optimization phase was re-trained on the small optimization sub-dataset (20,592 with the average activity of 0.32) to improve the biological activity of generated enhancers (**Fig. 1C**). The domain constraint was examined per epoch by calculating the MMD between the validation dataset and the generated dataset. The activity of generated enhancers was predicted by DeepSTARR. The generated sequences with activity higher than threshold were used to update the training dataset. The activity threshold started at 1.0 and was raised by 0.02 when the average activity of generated sequences was higher than the threshold for 5 times. More details are shown in https://github.com/chen-bioinfo/Enhancer-GAN.

### B. Capturing the general character of enhancers

In the first phase, we pre-trained WGAN to capture the general characteristics of real enhancers. We compared the effectiveness of pre-trained WGAN with the position-specific scoring matrix (PSSM) sampling method [12] and Gupta [6]. PSSM generated samples according to the natural enhancer base frequencies. The difference between our pre-trained model and Gupta is that we deprecate the strategy of Gumbel in the generator and focus more on the current information in every residual block. **Table I** shows the results of MMD in eight settings, including two embedding methods with four tokenization methods. The MMD value of our model is the closest to the positive control and away from the negative control. Although all of them have the same GC contents (0.4096) as that of real sequences (0.4099), the generated sequences from our model are closer to the real distribution compared to the others on MMD, which indicates our model can capture more information, such as nucleotide content and relation among nucleotides.

**Table 1:** The performance of our proposed model at first phase in terms of MMD based on Spectrum kernel embedding and DNABert embedding. The Positive Control simulates the best results between the generated distribution and the real one, while the Negative Control represents the worst results.

| Methods | Spectrum ($k$-mer) | | | | DNABert ($k$-mer) | | | |
| --- | --- | --- | --- | --- | --- | --- | --- | --- |
| | $k=3$ | $k=4$ | $k=5$ | $k=6$ | $k=3$ | $k=4$ | $k=5$ | $k=6$ |
| Positive Control | 0.038 | 0.038 | 0.033 | 0.025 | 0.030 | 0.020 | 0.018 | 0.020 |
| Negative Control | 0.341 | 0.327 | 0.251 | 0.168 | 0.264 | 0.151 | 0.126 | 0.134 |
| **Our** | **0.075** | **0.081** | **0.068** | **0.056** | **0.117** | **0.059** | **0.044** | **0.041** |
| PSSM | 0.225 | 0.227 | 0.181 | 0.126 | 0.146 | 0.082 | 0.065 | 0.074 |
| Gupta | 0.111 | 0.126 | 0.110 | 0.084 | 0.154 | 0.089 | 0.073 | 0.069 |

In addition, the t-SNE visualization (**Fig. 2**) between real enhancers and our generated enhancers with embeddings from DeepSTARR [3] shows that the generated enhancers occupy a similar data manifold as the real and enhancers with high activity appear in the same regions, which mean the generator can capture activity feature from raw sequence.

**Fig. 2:**
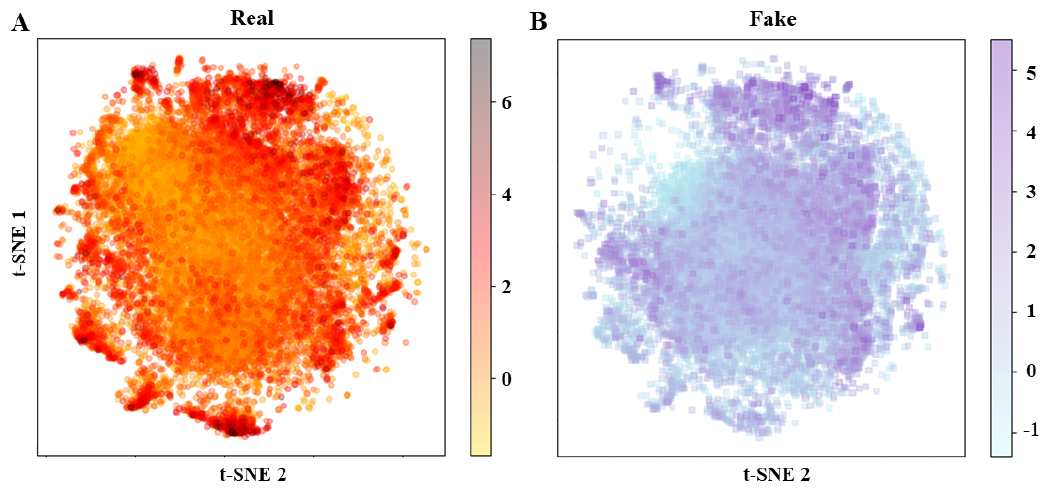
The t-SNE visualization between real samples and generated samples. The bar indicates the sequence activity.

### C. Improving sequence activity with feedback optimization

In the second phase, we added the feedback-loop mechanism into WGAN framework and re-trained on the feedback optimization sub-dataset to optimize the generator for high-activity enhancer generation.

#### Domain constraint

Because of the advantage of low-time cost, we employed the normalized Spectrum kernel with *mer* = 3 to evaluate the domain gap between the generated sequences and the validation dataset in the feedback loop. **Fig. 3** shows the impact of domain constraints with five constraint thresholds, *T*_*gap*_ = 0.2, 0.3, 0.4, 0.6, 1.0. When the constraint is loosened in the feedback mechanism, the generator can produce more generated samples with higher activity. However, loose constraints will introduce more noise of the external analyzer into the generator. With the another consideration of the MMD of 0.4603 between real enhancer sequences with low activity (less than -0.5) and high activity (higher than 3.0), we set the domain constraint gap to 0.3 in the following experiment to generate high-activity sequences while alleviating the noise from the external analyzer.

**Fig. 3:**
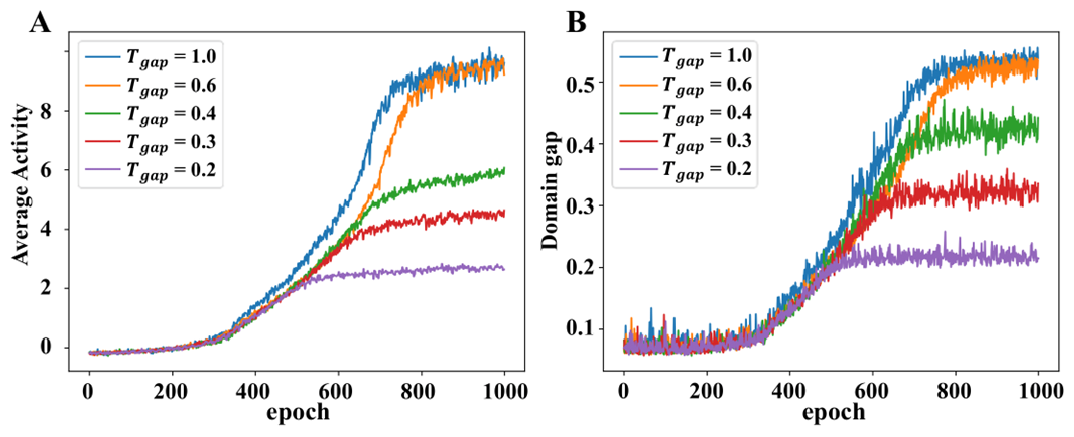
Impact of domain constraint for the generator during the re-training with feedback mechanism. (A) shows the impact for the average activity of the generated sequences, and (B) exhibits the value of MMD between generated sequences and validation dataset.

#### Curriculum learning

Given the average activity of 0.32 in the optimization sub-dataset, the cutoff started at 1.0 and then gradually increased during curriculum learning. When the average activity of generated samples exceeds cutoff for 5 times, cutoff was raised by 0.02. As shown in **Fig. 4A**, the WGAN makes little improvement for the activity of generated sequences. In contrast, the feedback mechanism significantly improves the activity of generated sequences. Moreover, Enhancer-GAN demonstrates a more rapid convergence compared to the feedback mechanism. **Fig. 4B-D** shows the activity distribution of generated sequences under three strategies in the optimization process. **Fig. 5A-C** shows the t-SNE analysis (based on the embedding from DeepSTARR) between generated sequences and real sequences in the epochs of 100, 300, and 500. We observed that generated sequences and real ones with the same activity are distributed in the same region, and there is an increasing proportion of high activity in generated sequences. In addition, the result from CD-HIT [24] shows the similarity between generated sequences in the 500-th epoch is less than 80%, which demonstrates their diversity.

**Fig. 4:**
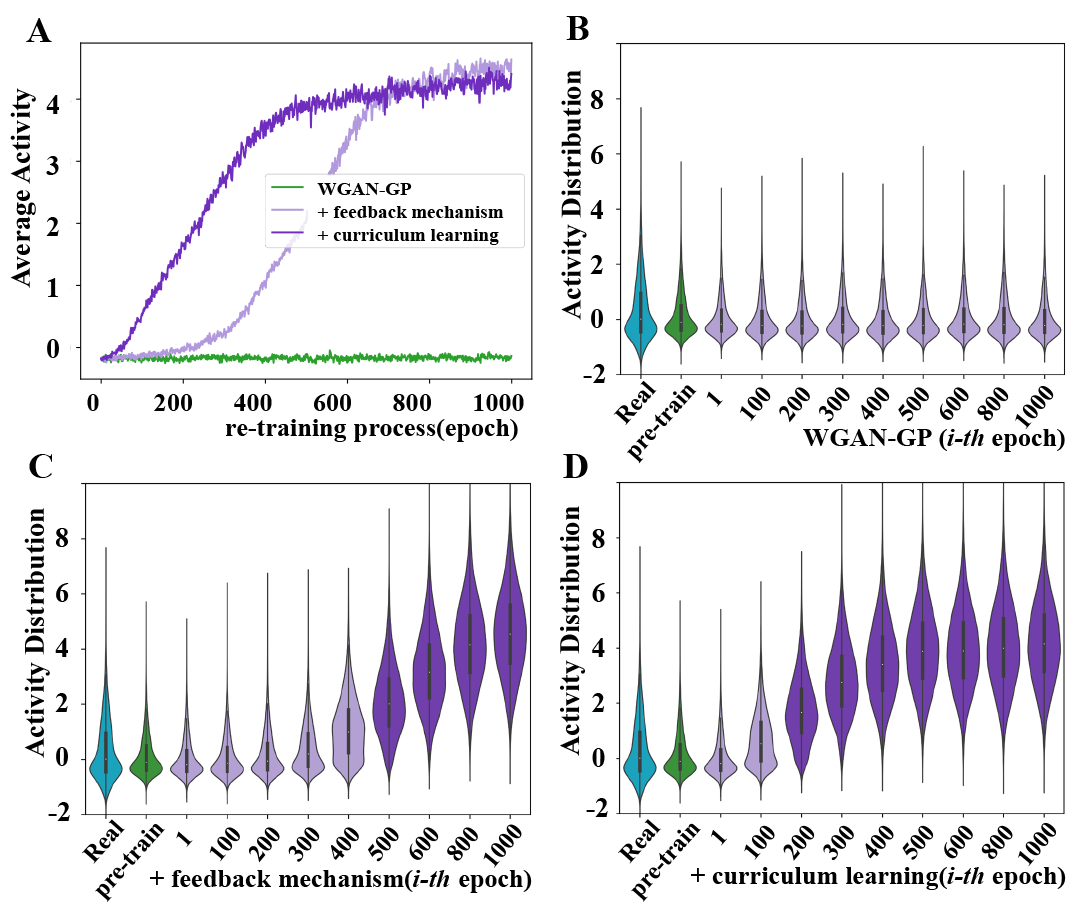
The activity distribution of generated sequences under three strategies in the optimization process. (A) The average activity of generated sequences in WGAN, feedback mechanism and feedback mechanism with curriculum learning (Enhancer-GAN), respectively. The activity distributions under these strategies during the optimization process are shown in (B) WGAN, (C) feedback mechanism and (D) Enhancer-GAN.

**Fig. 5:**
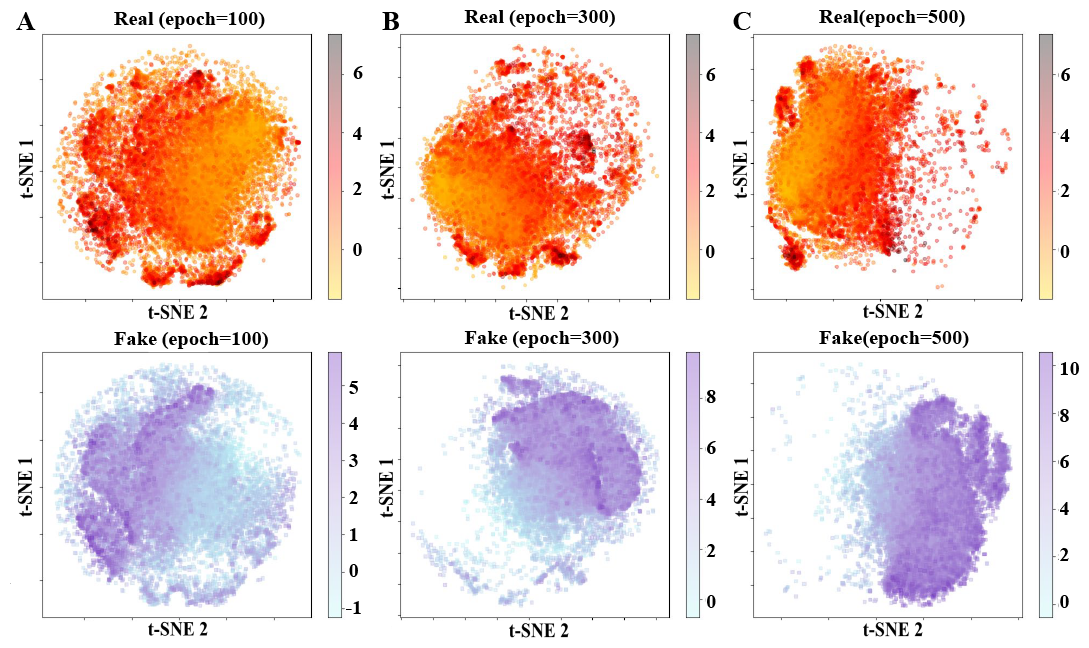
t-SNE visualization of real enhancers and generated enhancers by the generator with feedback mechanism and curriculum learning at epochs of 100, 300, 500 in the retraining process. At each epoch, 20,000 sequences were randomly generated by the generator.

### D. Enhancer motifs extraction via Enhancer-GAN

As mentioned in III-D, we sampled 20,000 input noise to generate 20,000 evolution sets and filtered them with the rule whether the sequence activity in each evolution set evolves in increasing order. At last, we collected 782 evolution sets, and divided them into three groups. Enhancers with activity less than 0.0 belong to low-activity group, larger than 3.0 belong to high-activity group, and others to mid-activity group. With the motif length of 8bp and P-value threshold of 0.005 in STREME, we extracted 10 fine motifs from the high-activity group and 10 flaw motifs from the low-activity group.

To figure out the relation between enhancer activity and its inner motifs, we swapped motifs between high-activity group and low-activity group, then observed the difference in activity (**Fig. 6**). At first, we replaced the flaw motif of ‘GGCTTATA’ contained in low-activity sequences with ten fine motifs and replaced ‘GGCTTATA’ contained in high-activity sequences with ten flaw motifs. In **Fig. 6A**, the replacements increase the activity of low-activity sequences at the average of 0.434, respectively. Secondly, **Fig. 6B** shows the results of the replacement of the fine motif of ‘GACTCACA’ contained in high-activity sequences to ten flaw motifs. The activity of these sequences decreases by the average of 1.039.

**Fig. 6:**
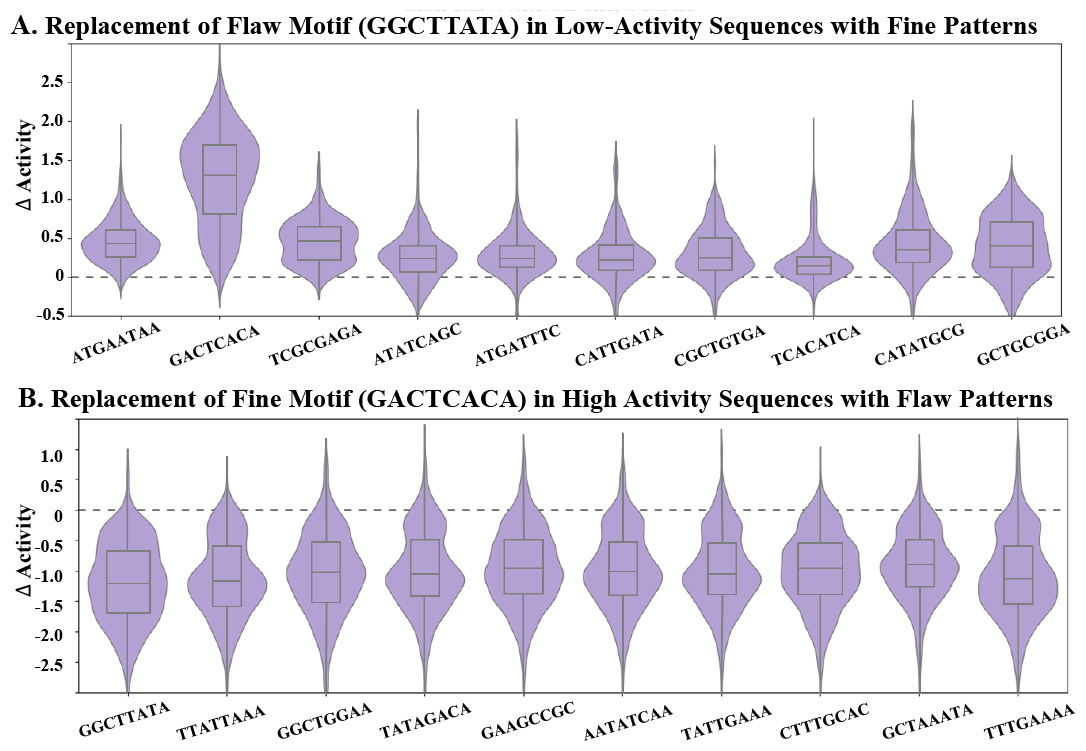
The activity changes caused by motif replacement among fine motifs and flaw motifs. (A) The activity of enhancers in low-activity is increased due to the replacement of the flaw motif (GGCTTATA) with 10 fine motifs. (B) The activity of enhancers in high-activity is decreased due to the replacement of fine motifs (GACTCACA) to 10 flaw motifs.

## V. Conclusion

In this paper, we developed an AI-driven enhancer design method, Enhancer-GAN, to produce enhancers with high activity. Enhancer-GAN is firstly pre-trained on a large enhancer dataset to learn the general grammar of real enhancers, then is optimized by feedback-loop with domain constraint and curriculum learning for high-activity enhancer generation. Therefore, the activity of generated enhancers slowly shifts over time. We have shown the robustness of domain constraint and curriculum learning to improve the activity of generated enhancers. Furthermore, we found 10 novel fine motifs extracted from generated high-activity enhancers, which demonstrates the promising ability to unveil the biological mechanism of enhancers.

In addition, Enhancer-GAN is not limited to generating Drosophila enhancer, but also human enhancer. In future work, we would also like to apply and further validate the currently proposed method on additional application areas, such as protein and drug design.

## VI. ACKNOWLEDGEMENT

This work is supported by the Natural Science Foundation of China (Grant No. 62102118), Project of Educational Commission of Guangdong Province of China (Grant No. 2021KQNCX274), the Shenzhen Colleges and Universities Stable Support Program (Grant No. GXWD20220811170504001)

